# Cryo-EM shows how dynactin recruits two dyneins for faster movement

**DOI:** 10.1101/183160

**Authors:** Linas Urnavicius, Clinton K. Lau, Mohamed M. Elshenawy, Edgar Morales-Rios, Carina Motz, Ahmet Yildiz, Andrew P. Carter

## Abstract

Dynein and its cofactor dynactin form a highly processive microtubule motor in the presence of an activating adaptor, such as BICD2. Different adaptors link dynein/dynactin to distinct cargos. Here we use electron microscopy (EM) and single molecule studies to show that adaptors can recruit a second dynein to dynactin. Whereas BICD2 is biased toward recruiting a single dynein, the adaptors BICDR1 and HOOK3 predominantly recruit two. We find that the shift toward a double dynein complex increases both force and speed. A 3.5 Å cryo-EM reconstruction of a dynein tail/dynactin/BICDR1 complex reveals how dynactin can act as a scaffold to coordinate two dyneins side by side. Our work provides a structural basis for how diverse adaptors recruit different numbers of dyneins and regulate the motile properties of the dynein/dynactin transport machine.

## Introduction

Human cytoplasmic dynein-1 (dynein) is the main transporter of cargos toward the minus ends of microtubules in eukaryotic cells (Roberts et al. 2013). The cargos it carries move over a range of different speeds (Klinman et al. 2016) and vary in size from large organelles (Burkhardt et al. 1997) to small, individual proteins (Ben-Yaakov et al. 2012). Dynein is converted into a highly processive motor by binding its cofactor dynactin and a cargo adaptor, such as BICD2 (Bicaudal D homolog 2) (McKenney et al. 2014; Schlager et al. 2014b). Mutations in any of the components of this dynein/dynactin transport complex are linked to neurodevelopmental and neurodegenerative disorders (Lipka et al. 2013). Dynein contains two motor domains held together by a tail region, whereas dynactin is built around a short actin-like filament, capped by the pointed-and barbed-end complexes and decorated with a large shoulder domain (Schlager et al. 2014b; Chowdhury et al. 2015; Urnavicius et al. 2015; Zhang et al. 2017). A previous 8 Å cryo-electron microscopy (cryo-EM) structure showed how a long coiled coil in BICD2 recruits dynein’s tail to dynactin’s filament (Urnavicius et al. 2015). A number of other adaptors have been identified that can activate dynein (McKenney et al. 2014; Olenick et al. 2016; Redwine et al. 2017) and link it to different cargos (Griffis et al. 2007; Horgan et al. 2010; Bielska et al. 2014; Zhang et al. 2014). These activating adaptors all contain long coiled coils and some of them share a limited number of short conserved motifs (Schlager et al. 2014a; Gama et al. 2017; Zheng 2017). However, in general the sequence similarity between them is low (Cianfrocco et al. 2015) making it unclear if they all engage dynein/dynactin in the same way. There is also evidence that certain adaptors, such as BICDR1 (Schlager et al. 2014a) (BICD related-1) and HOOK3 (McKenney et al. 2014; Olenick et al. 2016; Redwine et al. 2017), drive faster movement than BICD2, though the mechanism by which they achieve this is not understood.

## Dynactin can recruit two dyneins

To address these questions we first determined cryo-EM structures of two new dynein/dynactin/adaptor complexes. BICDR1, like BICD2, recruits dynein to Rab6 vesicles in cells (Schlager et al. 2010) but has yet to be shown to directly activate dynein/dynactin *in vitro*. HOOK3 is a well-established activating adaptor (McKenney et al. 2014; Olenick et al. 2016; Schroeder et al. 2016), which links dynein/dynactin complexes to early endosomes (Bielska et al. 2014; Zhang et al. 2014; Guo et al. 2016). We combined these adaptors with dynactin and a recombinant dynein tail complex, containing residues 1-1455 of the dynein heavy chain (HC) and its accessory chains. We determined ~7 Å resolution maps of both the dynein tail:dynactin:BICDR1 (TDR) and the dynein tail:dynactin:HOOK3 (TDH) complexes, which we compare to our previous structure of dynein tail:dynactin:BICD2 (TDB) (Urnavicius et al. 2015) (Figs. 1A, S1A-D, Table S1).

**Figure 1:**
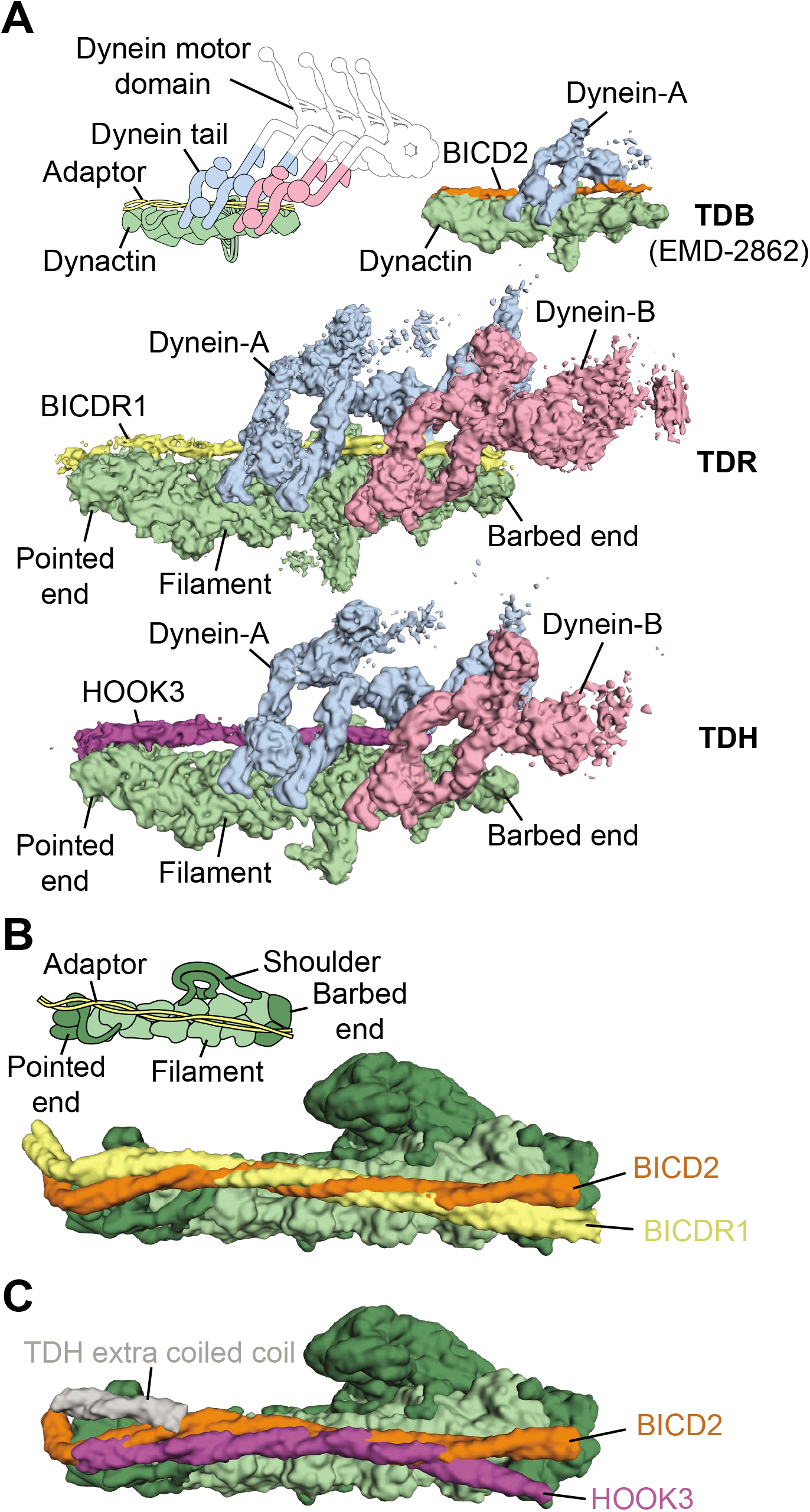
Dynactin can recruit two dyneins. **A.** Sub-7 Å cryo-EM density maps of the dynein tail:dynactin:BICDR1 (TDR) and tail:dynactin:HOOK3 (TDH) complexes, segmented and colored according to their components. tail:dynactin:BICD2 (TDB; EMD-2862) is included for comparison. **B.** Comparison of molecular models (surface representation) of BICDR1 with BICD2 coiled coils on dynactin show divergent contacts at the pointed and barbed ends. **C.** Comparison of HOOK3 and BICD2 coiled coils on dynactin’s surface.

The coiled coils of all three adaptors run along the length of the dynactin filament (Fig. 1A). However, in contrast to previous predictions (Zheng 2017) each adaptor makes remarkably different interactions (Fig. 1B). BICD2 and BICDR1, for example, diverge both in their path and relative rotation, especially toward the pointed and barbed ends of dynactin. HOOK3 follows yet another route over dynactin’s surface and also makes additional contacts with dynein/dynactin compared to BICDR1 (Figs. 1B, C, S1E, F). There is extra coiled-coil density at the dynactin pointed end (Fig. 1C) and extra globular density toward the barbed end (Fig. S1F). Whereas the identity of the second coiled coil is not clear, the globular density likely corresponds to the N-terminal Hook domain, which is required for HOOK3’s function as an activating adaptor (Olenick et al. 2016; Schroeder et al. 2016).

The most striking feature of the TDR and TDH structures is the presence of two dynein tails (Fig. 1A). The first dynein, which we call dynein-A, binds in an equivalent position to the single dynein tail in TDB (Urnavicius et al. 2015) and the full-length dynein in our dynein:dynactin:BICD2 (DDB) structure (Zhang et al. 2017). The second dynein, dynein-B, binds next to dynein-A close to the barbed end of dynactin. The dynein tails lie nearly parallel to each other in adjacent grooves along the dynactin filament.

## Adaptors determine dynein recruitment

We next determined whether BICD2, BICDR1 and HOOK3 recruit a different number of dyneins in moving dynein/dynactin complexes. We mixed tetramethylrhodamine (TMR)-labeled dynein and Alexa647-labeled dynein and used single molecule fluorescence microscopy to measure the frequency at which two dyes colocalize on microtubules (Fig. 2A, B). Dynein alone showed only 2.1±0.3% (s.e.m.) colocalization. In the presence of dynactin and BICD2, 13±1% of processive complexes were labeled with both dyes, significantly higher (P < 0.0001) than the dynein control. Using either BICDR1 or HOOK3 as an adaptor led to 31±2% and 34±1% colocalization respectively (Fig. 2B). After correction for complexes double-labeled with the same color (see Methods), we estimated that BICD2 results in 26% of complexes containing two dyneins, compared to 61% for BICDR1 and 67% for HOOK3. We conclude that motile complexes containing BICD2 can recruit either one or two dyneins, although the majority contain one. In contrast both BICDR1 and HOOK3 preferentially recruit two dyneins.

**Figure 2:**
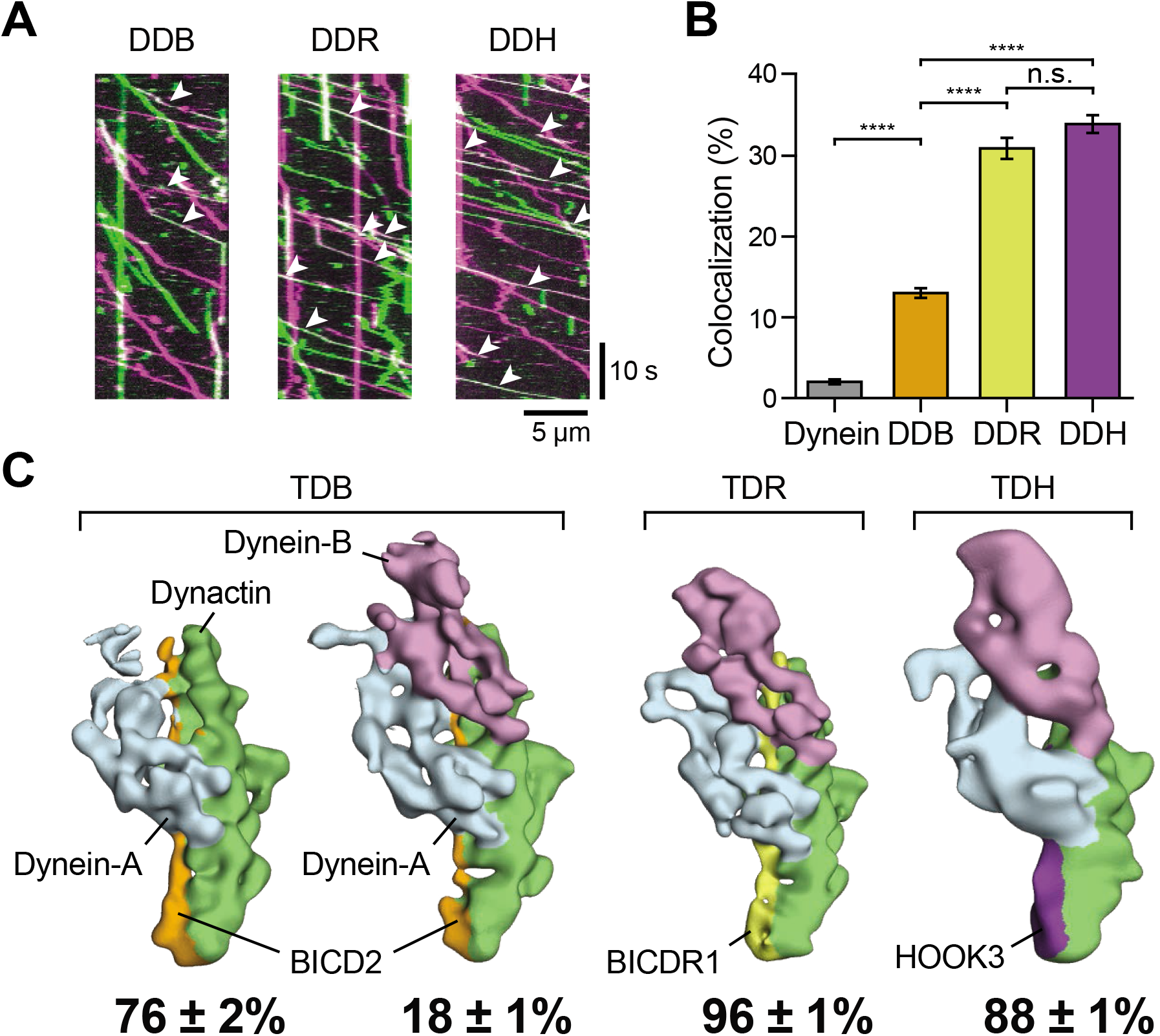
Adaptors recruit different number of dyneins to dynactin. **A.** Typical kymographs of dynein/dynactin assembled with BICD2 (DDB), BICDR1 (DDR) and HOOK3 (DDH) showing individual complexes containing signal from TMR-dynein (magenta), Alexa647-dynein (green) or both fluorescent dyneins (white). All colocalized events are marked with white arrowheads. **B.** The mean percentage (± s.e.m.) of complexes that contain both TMR-and Alexa647-dynein for each sample (N_Dynein_=7793, N_DDB_=3092, N_DDR_=3107, N_DDH_=3990; one-way ANOVA analysis; \*\*\*\**P* < 0.0001; n.s. - not significant). **C.** Representative 3D classes for one - and two-dynein complexes for TDB, TDR and TDH. Mean (± s.e.m.) fraction of particles for each class shown below (ambiguous classes are not shown).

Whereas this and previous studies (Schlager et al. 2014b,· Chowdhury et al. 2015; Urnavicius et al. 2015; Zhang et al. 2017) are consistent with BICD2 predominantly recruiting one dynein, its ability to recruit a second dynein was unanticipated. To obtain structural evidence to support this observation we applied a mixture of BICD2, dynein tail and dynactin onto grids for negative stain EM analysis (Fig. 2C). In agreement with our single molecule data, 3D classification of adaptor complexes showed that 18±1% of BICD2 complexes contained two dyneins, whereas 76±2% contained one, with the rest ambiguous. This ability of BICD2 to bind two dyneins also agrees with a cryo-electron tomography reconstruction of DDB, from brain lysate, bound to microtubules (G. Lander, personal communication). Negative stain EM of BICDR1 and HOOK3 complexes showed that 96±1% and 88±1% respectively contained two dyneins with the rest ambiguous (Fig. 2C). This suggests an even higher degree of second dynein recruitment than our single molecule data. Our combined data suggest that the number of dyneins bound to dynactin can be controlled by the identity of the adaptor.

## Two dyneins increase force and speed

We sought to understand how recruiting different numbers of motors affects the motile properties of the dynein/dynactin complex. We first used an optical trap to measure the stall force of DDB, dynein/dynactin/BICDR1 (DDR) and dynein/dynactin/HOOK3 (DDH) (Fig. 3A, B). Similar to our previous measurements (Belyy et al. 2016), the stall force of DDB is 3.7±0.2 pN, significantly lower (P < 0.0001) than the stall force of the plus end directed motor, kinesin-1 (5.7±0.2 pN) (Svoboda et al. 1993). In comparison, the stall force of DDR is 6.5±0.3 pN and DDH is 4.9±0.2 pN (Fig. 3B), suggesting that recruiting more dyneins to dynactin increases force production.

**Figure 3:**
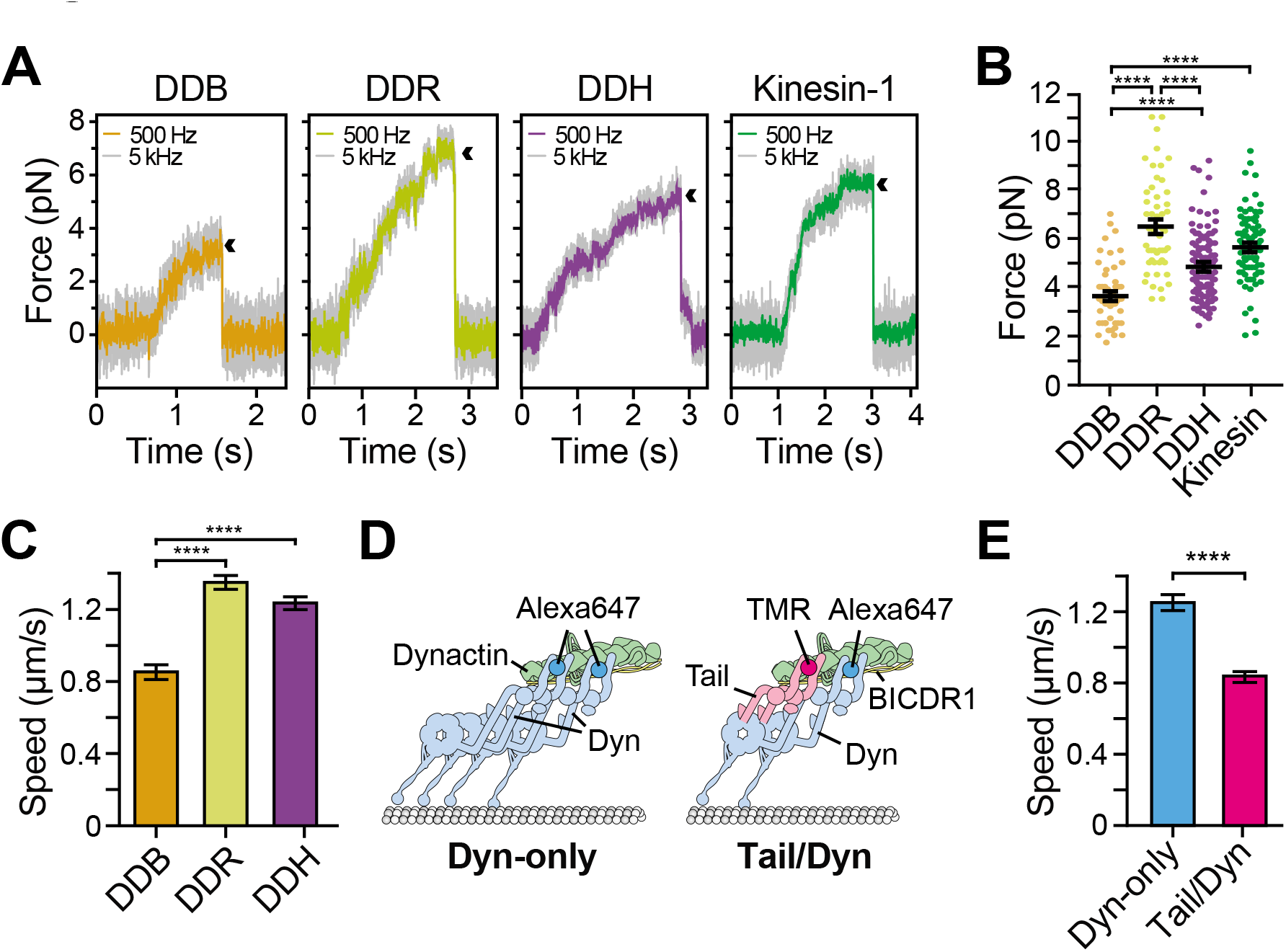
Two dyneins increase force and speed of dynein/dynactin. **A.** Traces showing typical stalls of beads driven by single DDB, DDR, DDH or human kinesin-1. Arrowheads denote the detachment of the motor from a microtubule after the stall. **B.** Scatter plots of optical trap data showing stall force distributions (N_DDB_=54, N_DDR_=53, N_DDH_=118, N_kinesin_=83). The horizontal line denotes the mean stall force. **C.** Mean speeds of DDB, DDR and DDH (N_DDB_=3343, N_ddr_=3162, N_ddh_=3744). **D.** Schematic depicting experimental design for TMR-tail/Alexa647-Dyn experiment. **E.** Dyn-only complexes move significantly faster than Tail/Dynein (NDyn-only= 1004, N_Taiı_/_Dyn_=939). In B, C and E error bars represent s.e.m.; \*\*\*\**P* < 0.0001 (ANOVA for B, C; unpaired t-test for E).

This agrees with previous reports that the stall force of a kinesin-or dynein-driven bead increases with the number of motors (Vershinin et al. 2007; Shubeita et al. 2008; Rai et al. 2013). However, more recent work with tighter control over motor copy number, using a DNA or protein scaffold, revealed that coupling multiple kinesins does not lead to a significant increase in stall force (Jamison et al. 2012; Furuta et al. 2013). The increased forces we observe for DDR and DDH may be explained by dynein motors resisting against microtubule detachment more effectively than kinesins under high backward loads (Jamison et al. 2012; Cleary et al. 2014; Dogan et al. 2015; Nicholas et al. 2015), enabling them to team up efficiently for maximum mechanical output (Rai et al. 2013; Rai et al. 2016). The difference in stall force between DDR and DDH, which both recruit two dyneins, suggests that other features of these specific dynein/dynactin complexes besides motor number can also fine tune force production.

The higher stall force of DDR also suggests it competes more efficiently with kinesin than DDB does. This may explain why neuronal overexpression of BICDR1, but not BICD2, counteracts kinesin-driven transport of Rab6 vesicles (Schlager et al. 2014a) and may be relevant to BICDR1’s biological role in opposing anterograde movement in early neuronal differentiation (Schlager et al. 2010). The ability of some adaptors to recruit multiple dyneins could also contribute to the clustering and pairing of dynein motors required to transport large cargos (Hendricks et al. 2012; Rai et al. 2016).

We next asked if recruiting more dyneins to dynactin had an effect on speed. Previous work on BICDR1 in cells (Schlager et al. 2014a) and HOOK3 *in vitro* (McKenney et al. 2014; Olenick et al. 2016; Schroeder et al. 2016; Redwine et al. 2017) showed complexes containing these adaptors move faster than those with BICD2. Our data raise the possibility these faster speeds are due to more complexes containing two dyneins. However, artificial tethering of processive dynein or kinesin motors had suggested motor number has little or no effect on velocity (Derr et al. 2012; Jamison et al. 2012; Furuta et al. 2013; Driller-Colangelo et al. 2016).

To determine whether motor number affects the speed of dynein/dynactin complexes, we first directly compared the speeds of all three adaptors in our *in vitro* motility assay (Figs. 3C, S2A, B). The average velocities of DDR (1.35±0.04 μm/s) and DDH (1.23±0.04 μm/s) were significantly faster than DDB (0.86±0.04 μm/s, *P* < 0.0001). To test whether this speed difference required the presence of two active dyneins we mixed Alexa647-labeled dynein with a TMR-labeled tail construct in the presence of BICDR1 and dynactin (Fig. 3D). We then compared the speeds of moving complexes containing only full length dynein (Dyn-only) with those that contained both dyes and hence one tail and one active dynein (Tail/Dyn). For this experiment we used a mutated version of full-length dynein that does not form the inhibited phi particle and so makes an efficient complex with dynactin (Zhang et al. 2017), similar to the dynein tail (Urnavicius et al. 2015). As expected, Dyn-only complexes moved at a similar speed (1.25±0.04 μm/s, Fig. 3E) to DDR formed with either wild-type dynein (1.35±0.04 μm/s, Figs. 3C, S2A) or the mutant dynein (1.22±0.05 μm/s, Fig. S2C). However, Tail/Dyn complexes moved significantly slower (0.84±0.03 μm/s, *P* < 0.0001, Figs. 3E, S2D). This suggests that the presence of a second dynein increases the velocity of dynein/dynactin complexes.

We propose that the increase in speed upon the recruitment of two dyneins is linked to the way in which dynactin recruits them side by side. This may restrict the inherent sideways and backwards movements of the motor domains (Reck-Peterson et al. 2006) to take a more direct and faster route along the microtubule. Intriguingly, the velocity of BICD2 complexes containing both fluorophores, and hence two dyneins (1.08±0.03 μm/s, Fig. S2E), was significantly faster than the average DDB velocity (P < 0.0001), but not as fast as DDR. This suggests that in addition to recruiting two dyneins, certain adaptors also affect speed through small differences in the way that they recruit the motors to dynactin.

## The 3.5 Å cryo-EM structure of dynein tail:dynactin:BICDR1

To understand how dynactin recruits two dyneins, we collected a further ten cryo-EM datasets of the TDR complex, which allowed us to determine its structure to an overall resolution of 3.5 Å (Fig. S3, Table S1). In this map the resolution of the dynactin filament extended to 3.4 Å, whereas that of the dynein tails varied from 3.7 Å to below 6 Å at the periphery (Fig. S3C). To improve the tail density, we identified a set of overlapping regions that moved as rigid blocks by 3D classification. We subtracted the signal for dynactin and dynein regions outside these blocks from the raw particles and performed focused 3D classification and refinement (Fig. S4). This process was repeated multiple times, improving the definition of the rigid blocks at each cycle. This led to a set of 3.4 Å maps that together cover the entire length of the dynein tail (Fig. S4, Table S1).

Previous low-resolution structures showed that the dynein tail comprises two heavy chains (HCs), each consisting of a series of helical bundles, that are held together by an N-terminal dimerization domain (NDD) (Urnavicius et al. 2015; Zhang et al. 2017). Each HC binds one intermediate chain (IC) and one light intermediate chain (LIC) (Schroeder et al. 2014; Chowdhury et al. 2015; Zhang et al. 2017). The ICs are held together by the dynein light chains, Robl, LC8 and Tctex, which bind to their extended N-terminal regions (Williams et al. 2007; Hall et al. 2010). Here, we use our high-resolution maps to build an atomic model of the dynein tail. We *de novo* traced helical bundles 1-6 of the HC and the WD40 domain of the IC (Figs. 4A, S5A, B, Table S1). We also placed helices for part of helical bundle 7 and rebuilt homology models for the LIC (Schroeder et al. 2014) and Robl (Hall et al. 2010) (Figs. 4A, S5A, C).

**Figure 4:**
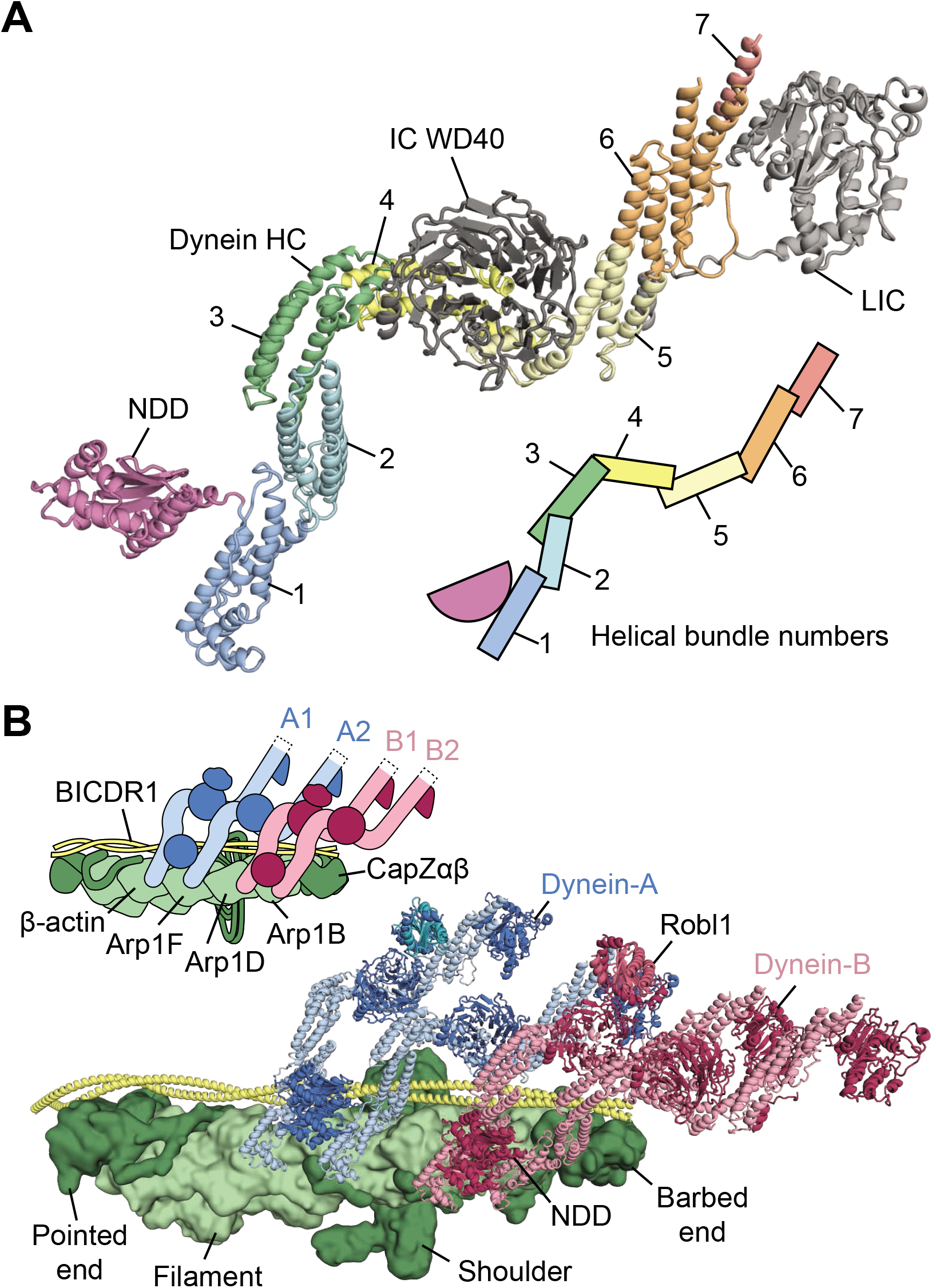
Structure of the dynein HC and architecture of the TDR complex. **A.** Consensus molecular model of one dynein heavy chain (HC), complete with intermediate chain (IC) and light intermediate chain (LIC). HC is colored according to helical bundle number. **B.** Assembled model of the TDR complex, showing the arrangement of the dynein-A (cartoon) and dynein-B (cartoon) on BICDR1 (cartoon) and dynactin (surface). The HC N-terminal dimerization domain (NDD) and dynein light chain Robl1 of dynein-B are labeled.

Our structure reveals that the IC WD40 domain makes extensive contacts to HC bundles 4-5, with a loop-helix from bundle 4 inserting into its central cavity (Fig. S6A). In contrast the LIC globular domain only interacts with two helices from bundle 6. Its tight binding to the HC (King et al. 2002) is a result of its N-and C-termini. These span out from the globular domain and form an integral part of HC bundles 5 and 7 respectively (Figs. S5C, S6B). It was previously observed that the Robl dimer, in complex with two IC N-terminal helices, binds an IC WD40 domain (Zhang et al. 2017). Intriguingly we find that this contact is solely mediated by the IC helices (Fig. S6C).

We finally assembled and refined a model of the whole TDR complex (Fig. 4B, Table S1) into our 3.5 Å map. We used our previous dynactin structure (Urnavicius et al. 2015) and a model for the BICDR1 coiled-coil region. For each dynein, we fit in two copies of HC/IC/LIC, one Robl dimer and a new 1.9 Å crystal structure of the human NDD (Fig. S6D, E, Table S2).

## The structural basis of two-dynein recruitment

Our TDR structure shows that the two dyneins, dynein-A and dynein-B, bind to the surface of the dynactin filament that is made up of the β-actin subunit and the three actin related protein 1 subunits Arp1F, Arp1D and Arp1B (Fig. 4B). The two dynein-A chains, A1 and A2, bind between the dynactin filament subunits β-actin/Arp1F and Arp1F/Arp1D respectively. The first dynein-B chain, B1, binds in the next groove between Arp1D and Arp1B. All three of these chains bind in a similar way, making contacts to both sides of their respective grooves. The precise interactions, though overlapping, are all slightly different (Fig. 5A). The second dynein-B chain, B2, binds between Arp1B and the barbed-end capping protein CapZβ. In order to bind this site, it is rotated by 90° along its long axis, relative to the other dynein chains.

**Figure 5:**
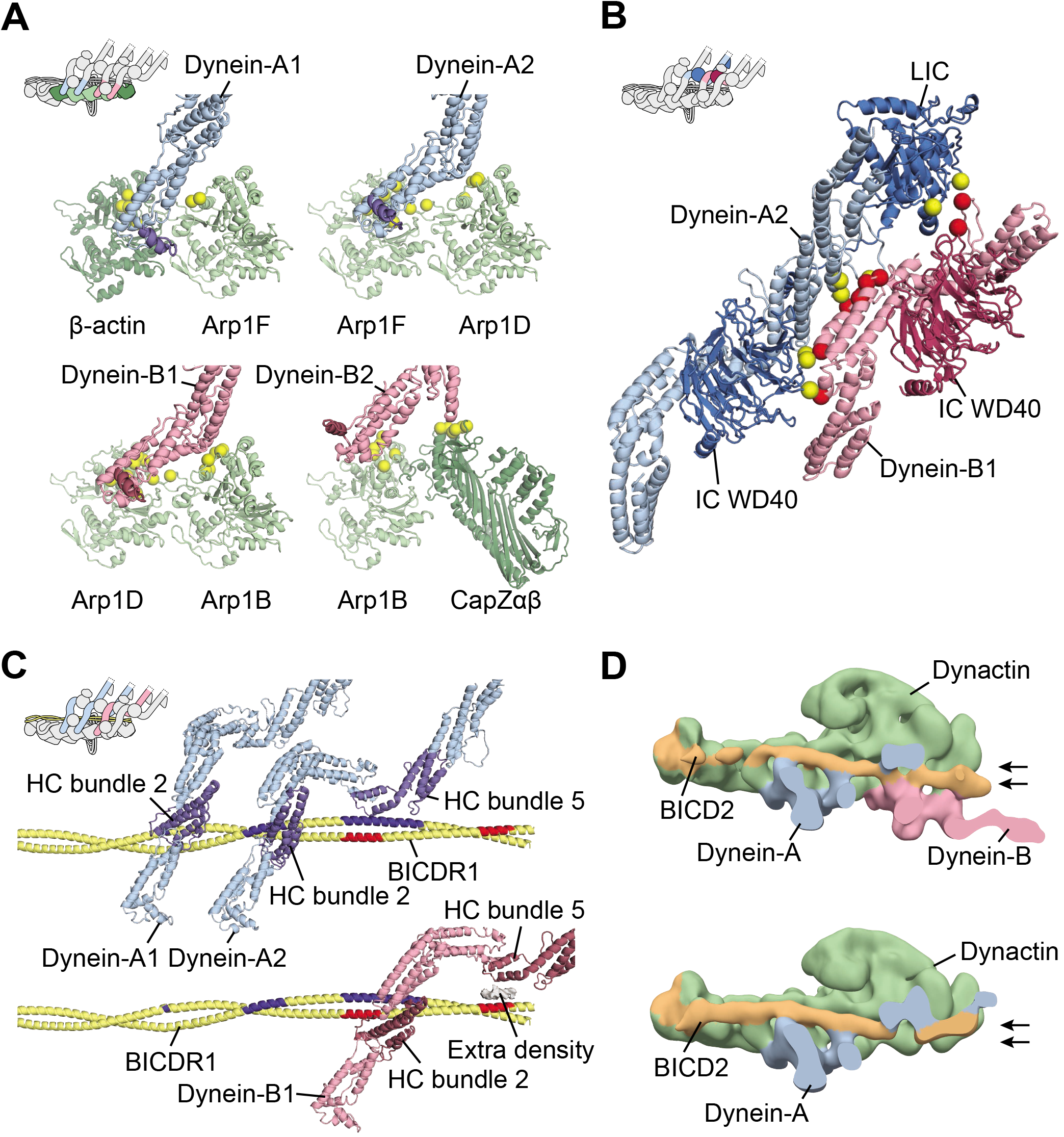
Interactions recruiting two dyneins to the TDR complex. **A.** Dynein HCs contacts dynactin similarly but using distinct residues (yellow spheres) in grooves along the dynactin filament (green). For each dynein HC, helix α6 is highlighted (dark blue or dark red). **B.** Dynein-A2 (blue cartoon, yellow spheres) makes extensive interactions with dynein-B1 (pink cartoon, red spheres). **C.** Both dynein-A chains and dynein-B1, but not dynein-B2, contact BICDR1. Interactions are shown on BICDR1 for dynein-A (dark blue) and dynein-B (red). Gray surface denotes the extra density between dynein-B1 and BICDR1. **D.** Negative stain EM reconstructions of DDB containing two dyneins (top) or dynein-A only (bottom), sliced to highlight the path of BICD2. Arrows depict the position of the BICD2 coiled coil at the barbed end of dynactin.

As well as binding to dynactin, the two dyneins make extensive interactions with each other. These consist of the IC WD40 domain of A2 binding the HC of B1; direct HC to HC interactions and contacts from the A2 LIC to both the HC and IC of dynein-B1 (Fig. 5B). These contacts contribute to a cascade of interactions between the 4 dynein chains (Fig. S7A) that involve direct contacts between the IC WD40 domain of each chain and the HC of its neighbor. This network of connections not only stabilizes the binding of the second dynein, but also ensures that all four HCs are held in a rigid orientation with respect to each other. This is likely to keep the dynein motor domains properly aligned with each other and may be important for the observed increase in speed when two dyneins are recruited to the dynactin scaffold.

Our structure also reveals the key role BICDR1 plays in recruiting two dyneins to dynactin. The first dynein, dynein-A, binds the adaptor in three places. Its A1 chain uses a single site on helical bundle 2 whereas the A2 chain binds via two sites (Fig. 5C). One of these also involves helical bundle 2, but is distinct from the interaction interface used by dynein-A1. The other involves a helix and loop on helical bundle 5. In comparison to dynein-A, recruitment of dynein-B depends only on its B1-chain as B2 makes no contacts with the adaptor. Dynein-B1 again uses sites on bundles 2 and 5. The first of these sites contacts the adaptor coiled coil opposite to where dynein-A2 binds (Fig. 5C). The second site does not directly contact the coiled coil, but instead touches density that packs against it (Figs. 5C, S7B, C). Although the identity of this region is uncertain, there is a weak density connecting it to helix 13 of LIC, suggesting it might correspond to the flexible C-terminus of the LIC (Fig. S7C). This region of LIC has been shown to interact with other activating adaptors (Schroeder et al. 2014; Schroeder et al. 2016).

## Adaptor position controls dynein number

To understand how different adaptors influence the number of dyneins bound, we also generated models of TDH and single-dynein-bound TDB using our low-resolution EM maps (Fig. 1A and EMDB-2860 (Urnavicius et al. 2015)). All three dynein/dynactin/adaptor complexes recruit dynein-A in a similar way despite the differences in the positions of the adaptors themselves (Fig. S8A). In TDR and TDH, dynein-B can bind because the BICDR1/HOOK3 N-termini follow downwards paths that are stabilized by interactions with CapZβ. In contrast, the BICD2 coiled coil is shifted upwards toward the shoulder domain to contact Arp1A in TDB (Figs. 1B, S8B). This prevents dynein-B1 binding by moving away and occluding its potential adaptor interaction site.

To address how BICD2 can also recruit two dyneins (Fig. 2), we asked what structural changes allow this adaptor to recruit a second dynein tail. We combined all of our replicate negative stain EM TDB datasets (Fig. 2C) to determine structures of sufficient quality to resolve the position of the coiled coil. We find that TDB with two dynein tails has the BICD2 is in a lower position (Fig. 5D). This is similar to that of BICDR1 and HOOK3, and different from the position in the TDB with only dynein-A bound (Fig. 5D). Our data therefore suggest that a switch in the position in the N-terminus of the adaptor is sufficient to account for the recruitment of dynein-B.

In conclusion, here we show that dynactin can act as a natural scaffold to line up two dyneins in close proximity. This arrangement results in a dynein/dynactin machine, which moves faster and can produce larger forces. Our observations provide a mechanism by which the cargo can control the output of the dynein/dynactin machine via the identity of its activating adaptor.

## Acknowledgements

We thank S. Scheres, X. Bai, K. Vinothkumar, A. Fitzpatrick, A. Brown, K. Zhang and R. Leiro for cryo-EM advice; S. Chen, G. McMullan, C. Savva, G. Cannone, J. Grimmett, and T. Darling for technical support at the MRC-LMB; S. Bullock for a SNAPf-dynein tail (1-1074-GST) construct; J.G. Shi for Sf9 cells; N. Barry and H.T. Hoang for microscopy advice; M. Yu for crystallography support and the European Synchrotron Radiation Facility (beamline ID29) for data collection; T. Croll for discussions on model building; S. Leech for fresh pig brains; S. Bullock, L. Passmore, S. Lacey, H. Foster, and J. Carter for comments on the manuscript; D. Grotjahn, S. Chowdhury and G. Lander for discussions. This work was funded by grants from the Wellcome Trust (WT100387) and the Medical Research Council, UK (MC_UP_A025_1011) to A.P.C., and the NIH (GM094522), and NSF (MCB-1055017, MCB-1617028) to A.Y.

## Author Contributions

L.U. performed all cryo-EM work on TDR and devised the strategy for high-resolution structure determination. C.K.L. determined the cryo-EM structure of TDH. L.U., C.K.L., M.M.E and A.P.C. measured and analyzed velocity and colocalization. L.U. carried out negative stain EM studies. M.M.E. and A.Y. performed optical trapping. E.M-R determined the X-ray structure of NDD. C.M. made the 1-1455 dynein tail construct. A.P.C., L.U. and C.K.L built and refined the TDR model and prepared the manuscript. A.P.C. initiated and supervised the project.

## Data availability

Cryo-EM maps were deposited in the Electron Microscopy Data Bank with accession IDs EMD-XXXX, EMD-XXXX and EMD-XXXX. Model coordinates were submitted to the Protein Data Bank (accession IDs XXXX, XXXX, XXXX). All data from this manuscript are available from the corresponding author.

## References

Belyy V. et al. 2016. The Mammalian Dynein–dynactin Complex Is a Strong Opponent to Kinesin in a Tug-of-War Competition. Nature Cell Biology 18(9): 1018–24.

Ben-Yaakov K. et al. 2012. Axonal Transcription Factors Signal Retrogradely in Lesioned Peripheral Nerve. The EMBO journal 31(6): 1350–63.

Bielska E. et al. 2014. Hook Is an Adapter That Coordinates Kinesin-3 and Dynein Cargo Attachment on Early Endosomes. Journal of Cell Biology 204(6): 989–1007.

Burkhardt, J. K., Echeverri, C. J., Nilsson, T., and Vallee, R. B. 1997. Overexpression of the Dynamitin (p50) Subunit of the Dynactin Complex Disrupts Dynein-Dependent Maintenance of Membrane Organelle Distribution. Journal of Cell Biology 139(2): 469–84.

Chowdhury, S., Ketcham, S. A., Schroer, T. A., and Lander, G. C. 2015. Structural Organization of the Dynein-Dynactin Complex Bound to Microtubules. Nature Structural & Molecular Biology 22(4): 345–47.

Cianfrocco, M. A., DeSantis, M. E., Leschziner, A. E., and Reck-Peterson, S. L. 2015. Mechanism and Regulation of Cytoplasmic Dynein. Annual Review of Cell and Developmental Biology 31(1): 83108.

Cleary F. B. et al. 2014. Tension on the Linker Gates the ATP-Dependent Release of Dynein from Microtubules. Nature Communications 5: 4587.

Derr N. D. et al. 2012. Tug-of-War in Motor Protein Ensembles Revealed with a Programmable DNA Origami Scaffold. Science 338(6107): 662–65.

Dogan M. Y. et al. 2015. Kinesin’s Front Head Is Gated by the Backward Orientation of Its Neck Linker. Cell Reports 10(12): 1968–74.

Driller-Colangelo, A. R., Chau, K. W. L., Morgan, J. M., and Derr, N. D. 2016. Cargo Rigidity Affects the Sensitivity of Dynein Ensembles to Individual Motor Pausing. Cytoskeleton 73(12): 693–702.

Furuta K. et al. 2013. Measuring Collective Transport by Defined Numbers of Processive and Nonprocessive Kinesin Motors. Proceedings of the National Academy of Sciences of the United States of America 110(2): 501–6.

Gama J. B. et al. 2017. Molecular Mechanism of Dynein Recruitment to Kinetochores by the Rod-Zw10-Zwilch Complex and Spindly. Journal of Cell Biology 216(4): 943–60.

Griffis, E. R., Stuurman, N., and Vale, R. D. 2007. Spindly, a Novel Protein Essential for Silencing the Spindle Assembly Checkpoint, Recruits Dynein to the Kinetochore. The Journal of Cell Biology 177(6): 1005–15.

Guo, X., Farías, G. G., Mattera, R., and Bonifacino, J. S. 2016. Rab5 and Its Effector FHF Contribute to Neuronal Polarity through Dynein-Dependent Retrieval of Somatodendritic Proteins from the Axon. Proceedings of the National Academy of Sciences 113(36): E5318–27.

Hall, J., Song, Y., Karplus, P. A., and Barbar, E. 2010. The Crystal Structure of Dynein Intermediate Chain-Light Chain Roadblock Complex Gives New Insights into Dynein Assembly. Journal of Biological Chemistry 285(29): 22566–75.

Hendricks, A. G., Holzbaur, E. L. F., and Goldman, Y. E. 2012. Force Measurements on Cargoes in Living Cells Reveal Collective Dynamics of Microtubule Motors. Proceedings of the National Academy of Sciences of the United States of America 109(45): 18447–52.

Horgan C. P. et al. 2010. Rab11-FIP3 Links the Rab11 GTPase and Cytoplasmic Dynein to Mediate Transport to the Endosomal-Recycling Compartment. Journal of Cell Science 123(2): 181–91.

Jamison, D. K., Driver, J. W., and Diehl, M. R. 2012. Cooperative Responses of Multiple Kinesins to Variable and Constant Loads. Journal of Biological Chemistry 287(5): 3357–65.

King, S. J., Bonilla, M., Rodgers, M. E., and Schroer, T. A. 2002. Subunit Organization in Cytoplasmic Dynein Subcomplexes. Protein science: a publication of the Protein Society 11(5): 1239–50.

Klinman, E., and Holzbaur, E. L. F. 2016. Comparative Analysis of Axonal Transport Markers in Primary Mammalian Neurons. In Methods in Cell Biology, 409–24.

Lipka, J., Kuijpers, M., Jaworski, J., and Hoogenraad, C. C. C. 2013. Mutations in Cytoplasmic Dynein and Its Regulators Cause Malformations of Cortical Development and Neurodegenerative Diseases. Biochemical Society Transactions 41(6): 1605–12.

McKenney R. J. et al. 2014. Activation of Cytoplasmic Dynein Motility by Dynactin-Cargo Adapter Complexes. Science 345(6194): 337–41.

Nicholas M. P. et al. 2015. Cytoplasmic Dynein Regulates Its Attachment to Microtubules via Nucleotide State-Switched Mechanosensing at Multiple AAA Domains. Proceedings of the National Academy of Sciences of the United States of America 112(20): 6371–76.

Olenick M. A. et al. 2016. Hook Adaptors Induce Unidirectional Processive Motility by Enhancing the Dynein-Dynactin Interaction. Journal of Biological Chemistry 291(35): 18239–51.

Rai A. et al. 2016. Dynein Clusters into Lipid Microdomains on Phagosomes to Drive Rapid Transport toward Lysosomes. Cell 164(4): 722–34.

Rai A. K. et al. 2013. Molecular Adaptations Allow Dynein to Generate Large Collective Forces inside Cells. Cell 152(1-2): 172–82.

Reck-Peterson S. L. et al. 2006. Single-Molecule Analysis of Dynein Processivity and Stepping Behavior. Cell 126(2): 335–48.

Redwine W. B. et al. 2017. The Human Cytoplasmic Dynein Interactome Reveals Novel Activators of Motility. eLife 6.

Roberts A. J. et al. 2013. Functions and Mechanics of Dynein Motor Proteins. Nature reviews. Molecular cell biology 14(11): 713–26.

Schlager M. A. et al. 2010. Pericentrosomal Targeting of Rab6 Secretory Vesicles by Bicaudal-D-Related Protein 1 (BICDR-1) Regulates Neuritogenesis. The EMBO journal 29(10): 1637–51.

Schlager M. A. et al. 2014a. Bicaudal D Family Adaptor Proteins Control the Velocity of Dynein-Based Movements. Cell Reports 8(5): 1248–56.

Schlager M. A. et al. 2014b. In Vitro Reconstitution of a Highly Processive Recombinant Human Dynein Complex. EMBO Journal 33(17): 1855–68.

Schroeder, C. M., Ostrem, J. M. L., Hertz, N. T., and Vale, R. D. 2014. A Ras-like Domain in the Light Intermediate Chain Bridges the Dynein Motor to a Cargo-Binding Region. eLife 3.

Schroeder, C. M., and Vale, R. D. 2016. Assembly and Activation of Dynein-Dynactin by the Cargo Adaptor Protein Hook3. Journal of Cell Biology 214(3): 309–18.

Shubeita G. T. et al. 2008. Consequences of Motor Copy Number on the Intracellular Transport of Kinesin-1-Driven Lipid Droplets. Cell 135(6): 1098–1107.

Svoboda, K., Schmidt, C. F., Schnapp, B. J., and Block, S. M. 1993. Direct Observation of Kinesin Stepping by Optical Trapping Interferometry. Nature 365(6448): 721–27.

Urnavicius L. et al. 2015. The Structure of the Dynactin Complex and Its Interaction with Dynein. Science 347(6229): 1441–46.

Vershinin M. et al. 2007. Multiple-Motor Based Transport and Its Regulation by Tau. Proceedings of the National Academy of Sciences of the United States of America 104(1): 87–92.

Williams J. C. et al. 2007. Structural and Thermodynamic Characterization of a Cytoplasmic Dynein Light Chain-Intermediate Chain Complex. Proceedings of the National Academy of Sciences, USA 104(24): 10028–33.

Zhang J. et al. 2014. HookA Is a Novel Dynein-early Endosome Linker Critical for Cargo Movement in Vivo. The Journal of Cell Biology 204(6).

Zhang K. et al. 2017. Cryo-EM Reveals How Human Cytoplasmic Dynein Is Auto-Inhibited and Activated. Cell 169(7): 1303–14.

Zheng W. 2017. Probing the Energetics of Dynactin Filament Assembly and the Binding of Cargo Adaptor Proteins Using Molecular Dynamics Simulation and Electrostatics-Based Structural Modeling. Biochemistry 56(1): 313–23.

